# Network-based approaches elucidate differences within APOBEC and clock-like signatures in breast cancer

**DOI:** 10.1101/568568

**Authors:** Yoo-Ah Kim, Damian Wojtowicz, Rebecca Sarto Basso, Itay Sason, Welles Robinson, Dorit S. Hochbaum, Mark D.M. Leiserson, Roded Sharan, Fabio Vandin, Teresa M. Przytycka

## Abstract

Studies of cancer mutations typically focus on identifying cancer driving mutations. However, in addition to the mutations that confer a growth advantage, cancer genomes accumulate a large number of passenger somatic mutations resulting from normal DNA damage and repair processes as well as mutations triggered by carcinogenic exposures or cancer related aberrations of DNA maintenance machinery. These mutagenic processes often produce characteristic mutational patterns called mutational signatures. Understanding the etiology of the mutational signatures shaping a cancer genome is an important step towards understanding tumorigenesis. Considering mutational signatures as phenotypes, we asked two complementary questions (i) what are functional pathways whose gene *expression* profiles are associated with mutational signatures, and (ii) what are *mutated pathways* (if any) that might underlie specific mutational signatures? We have been able to identify pathways associated with mutational signatures on both expression and mutation levels. In particular, our analysis provides novel insights into mutagenic processes in breast cancer by capturing important differences in the etiology of different APOBEC related signatures and the two clock-like signatures. These results are important for understanding mutagenic processes in cancer and for developing personalized drug therapies.

## 1 Introduction

Cancer genomes accumulate a high number of mutations, only a small portion of which are cancer driving mutations. Most of such mutations are passenger somatic mutations, not directly contributing to cancer development. Analyses of large scale cancer genome data revealed that these passenger mutations often exhibit characteristic mutational patterns called “mutational signatures” [1]. Importantly, these characteristic mutational signatures are often linked to specific mutagenic processes, making it possible to infer which mutagenic processes have been active in the given patient. This information often provides important clues about the nature of the diseases. For example, the presence of specific signatures associated with homologous recombination repair deficiency (HRD) can help identify patients who can benefit from PARP inhibitor treatment [2]. With the increased interest in the information on mutagenic processes acting on cancer genomes, several computational approaches have been developed to define mutational signatures in cancer [1, 3, 4, 5, 6, 7], to identify patients whose genome contains given signatures [6, 7, 8], and to map patient mutations to these signatures [9].

Despite the importance of understanding cancer mutational signatures, the etiology of many signatures is still not fully understood. It is believed that mutational signatures may arise not only as a result from exogenous carcinogenic exposures (e.g., smoking, UV exposures) but also due to endogenous causes (e.g., HRD signature mentioned above). That is, human genomes are protected by multiple DNA maintenance and repair mechanisms in the presence of various types of DNA damage but aberrations or other malfunctions in such mechanisms can leave errors not repaired, generating specific patterns of mutations [10].

From the perspective of individual patients, it is important to determine mutational signatures imprinted on each patient’s genome and the strength of the (sometimes unknown) mutagenic processes underlining the signatures. Signature strength can be measured by the number of mutations that are attributed to the given signature and thus can be considered as a continuous phenotype. With this view in mind, we investigate the relation of this phenotype with other biological properties of cancer patients. In this study, we focus on the relation of mutational signature strength with gene expression in biological processes and gene alteration in subnetworks.

The hypothesis that mutational signatures can be related to aberrant gene expression or alterations in DNA repair genes is well supported. For example, the deactivation of MUTYH gene in cancer patients is associated with a specific mutational signature [10, 11]. Previous studies identified correlations between several mutational signatures and some cancer drivers and acknowledged that the cause-effect relation between signatures and cancer drivers can be in either direction [12]. On the other hand, like many other cancer phenotypes, the causes of mutational signatures can be heterogeneous and the same signature can arise due to different causes. For example, the MUTYH signature mentioned above is known to be caused by inactivation of the MUTYH gene [13] but can also occur in cancers that do not harbor this aberration. With the observation that different mutations in functionally related genes can lead to the same cancer phenotype [14, 15, 16], cancer phenotypes are increasingly considered in the context of genetically dysregulated pathways rather than in the context of individual genes [17, 18, 19, 20, 21, 22]. Hence, we postulated that identifying mutated subnetworks and differentially expressed gene groups that are associated with mutational signatures can provide new insights on the etiology of mutational signatures.

In this study, we focused on mutational signatures in breast cancer, for which a large data set is available, including whole genome mutation profiles as well as expression data [23]. The mutagenic landscape of this cancer type is complex and is yet to be fully understood. For example, previously defined *COSMIC* signatures present in breast cancer [23] include two signatures (Signatures 1 and 5) as age related (clock-like) and two signatures associated with the activities of APOBEC enzyme (Signatures 2 and 13). The mechanisms underlying the differences between two distinct signatures with similar etiology are not fully understood.

The clock-like signatures (COSMIC Signatures 1 and 5) have been found correlated with the age of patients but the strengths of correlation differ between the two signatures and vary across different cancer types [24]. Signature 1 is considered to arise from an endogenous mutational process initiated by spontaneous deamination of 5-methylcytosine while the etiology of Signature 5 is less understood. Therefore, it is important to understand what processes, other than patient’s age, contribute to each of these signatures.

APOBEC signatures have been the subject of particular attention [25, 26, 27, 28, 29, 30, 31, 32, 33]. The proteins encoded by APOBEC gene family (known to be involved in immune response) deaminate cytosines in single-stranded DNA (ssDNA). Such deamination, if not properly repaired, can lead to C>T (Signature 2) or C>G (signature 13) mutations depending on how the resulting lesion is repaired or bypassed during the replication [34]. Thus the final imprint of APOBEC related mutations on the genome depends on several factors: expression level of APOBEC genes, the amount of accessible ssDNA, and the lesion bypass mechanism. In particular, clustered APOBEC-induced mutations (*kataegis*) in breast cancer are assumed to be a result of the mutation opportunity offered by single-stranded DNA during repair of double-stranded breaks (DSBs). However, ssDNA regions can also emerge for other reasons such as topological stress. Thus, although several aspects contributing to the APOBEC signatures have been known for some time, we are yet to uncover the full complexity of the APOBEC derived signatures.

To address these challenges, we took two complementary pathway based approaches: one focused on gene modules whose expression correlates with signature strength and the second based on the identification of subnetworks of genes whose alterations are associated with mutational signatures. Using our network/module based approaches, we were able to capture important differences in the etiology of different APOBEC related signatures and clock-like signatures in breast cancer.

Our study provides several new insights on the mutagenic processes in breast cancer including: (i) association of the NER pathway and oxidation processes with the strength of clock-like Signature 5 (ii) differences between the two clock-like signatures with respect to their associations with cell cycle (iii) differences in mutated subnetworks associated with different signatures including APOBEC related signatures. We demonstrate that our findings are consistent with the results from recent studies and provide additional insights that are important for understanding mutagenic processes in cancer and developing anti-cancer drugs.

## 2 Analysis Overview

In this study we consider mutational signatures in cancer patients and attempt to identify genes and pathways whose expression and/or genetic alterations are potentially causative of differences in mutational signature strength. Since recent studies revealed that mutations occurring in close proximity to each other, referred to here as cloud mutations, have distinct properties from dispersed mutations [35, 9], we additionally sub-divided all mutations (and subsequently their attributed signatures) into two groups – close-by **<u>C</u>**loud mutations and **<u>D</u>**ispersed mutations. We used only sufficiently abundant mutational signatures for cloud or dispersed mutations in the cohort of 560 breast cancer genomes [23], as inferred by a recently developed HMM based approach – SIGMA [9]. Consequently, we considered the 10 different phenotype profiles for Signatures 1D, 2C/D, 3C/D, 5D, 8C/D and 13C/D, where the numbering refers to the COSMIC signature index and C/D denotes close-by cloud and dispersed signatures. For details, see Materials and Methods section.

In the first part of the analysis, we looked for the genes whose expression levels are significantly correlated with mutational signature strength (Fig 1A,B). Specifically, we first selected genes exhibiting significant correlation with at least one mutational signature by computing the correlation coefficient of the expression profile and mutation counts for each pair of genes and signatures. The selected genes were clustered based on their expression correlation patterns across 10 mutational signatures. The enrichment analysis of each gene cluster revealed that different combinations of cellular processes are associated with different mutational signatures.

**Figure 1:**
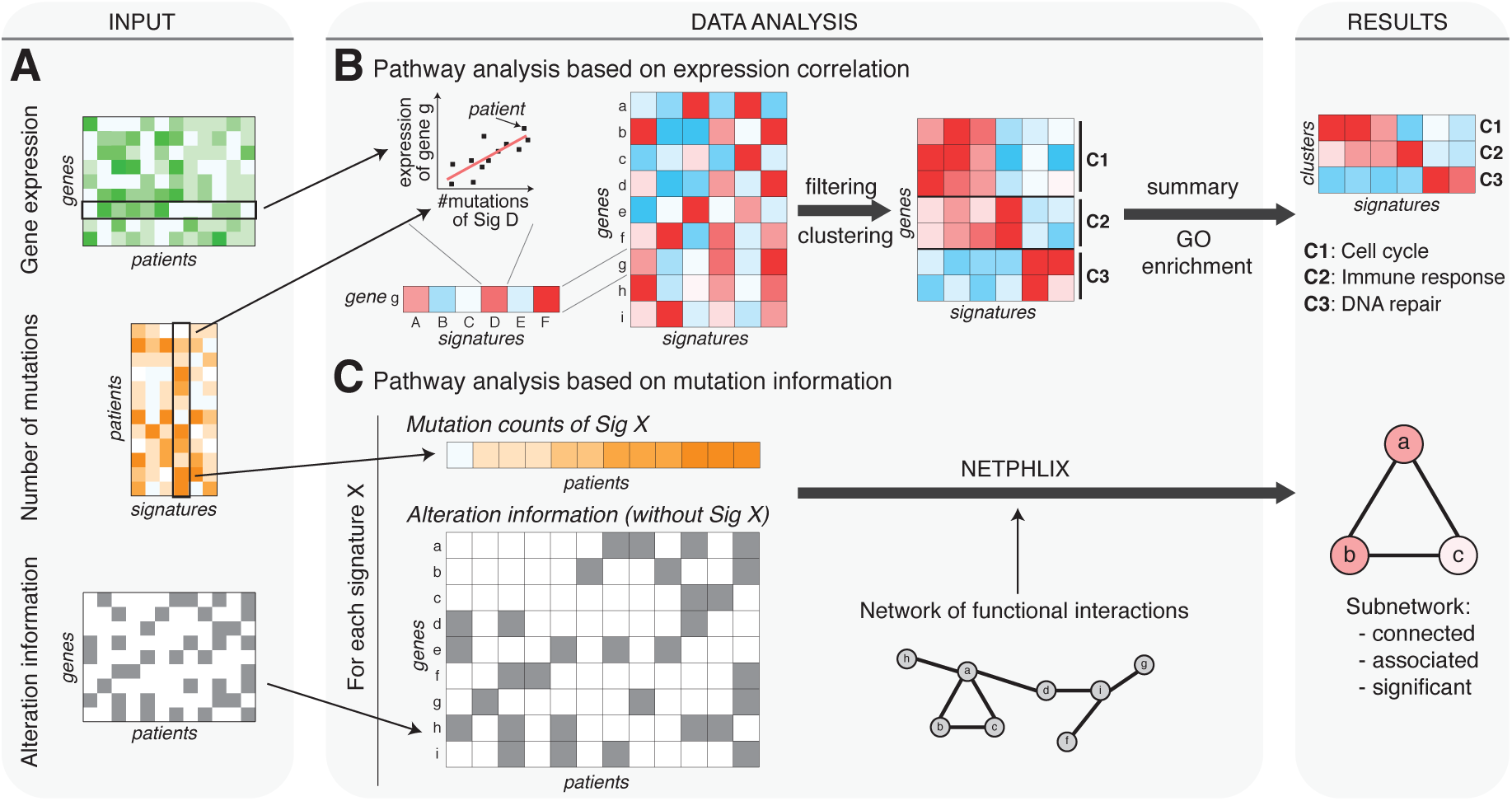
Overview of the study. **(A)** The input data for this study consist of gene expression, mutational signature frequency, and gene alteration across a number of cancer patients. **(B)** The functional pathways whose gene expression levels are associated with mutational signatures were found by computing correlations between expression levels of all genes and signature mutation counts, filtering out weak correlations, clustering expression correlation profiles, and performing GO enrichment analysis of the identified clusters. **(C)** The pathways whose gene alterations are associated with mutational signatures were found by applying NETPHIX to the signature mutation counts, gene-patient alteration matrix and a known functional interactions network.

**Figure 2:**
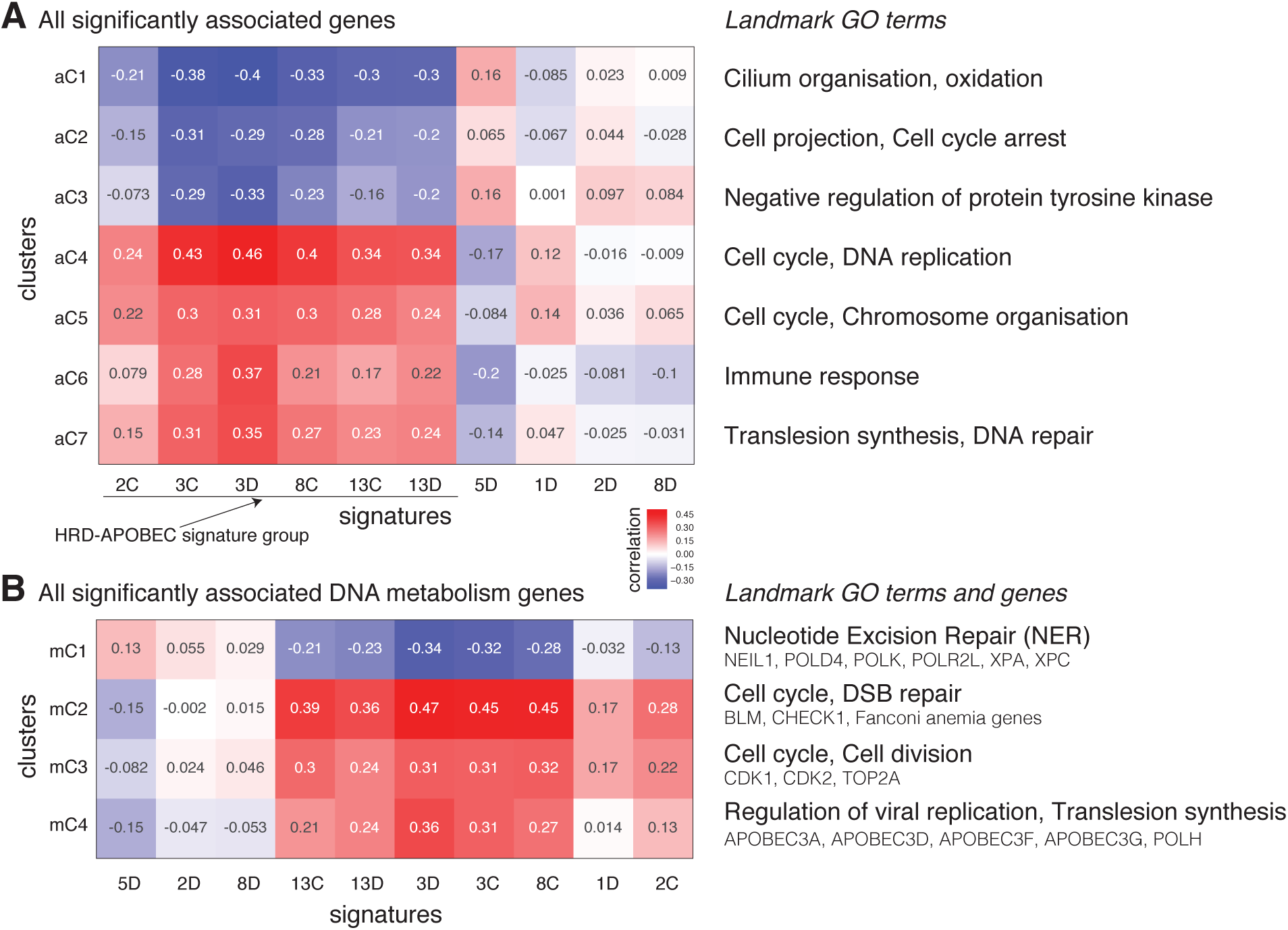
Gene expression correlation modules. **(A)** All genes significantly correlated with at least one signature. **(B)** DNA metabolic process genes, based on Gene Ontology (GO), significantly correlated with at least one signature. For both (A and B), we show a heatmap of mean expression correlation for each cluster and signature (left) and representative GO terms enriched in cluster genes (right). For the DNA metabolic process, we also show representative genes for each cluster. The list of GO enrichment terms is provided in the Supplementary Materials (Tables S1 and S2).

The second part of the analysis involves uncovering subnetworks of genes whose alterations are associated with mutational signature strength (Fig 1A,C). We hypothesize that a certain mutational signature can arise when a related pathway (e.g. DNA damage repair mechanism) is dysregulated. Due to the complex nature of cancer driving mutations, we adapted the NETPHIX method – a recently developed network based method to identify mutated subnetworks associated with continuous phenotypes [36]– to identify such pathways. In this analysis, we consider the mutation count of a mutational signature in a whole cancer genome to be the phenotype and aim to identify a subnetwork of genes whose alterations are associated with the phenotype. Importantly, when assessing association between gene level alterations and a mutational signature, the mutations attributed to the given mutational signature were not incorporated into the alteration information (Fig 1C; see Materials and Methods) in order to increase the likelihood of uncovered subnetworks being drivers of the signatures rather than their effect.

## 3 Results

### 3.1 Expression analysis to identify biological processes associated with mutational signatures

In order to identify biological processes associated with individual signatures, we clustered gene expression-signature correlation profiles as described in the Methods section. To obtain a bird’s eye view, we first used all genes whose expression is correlated with at least one signature (Figure 2A; see Materials and Methods). Next, to obtain a finer scale expression modules related to DNA repair, we zoomed in on genes involved in Gene Ontology DNA metabolic process (Figure 2B).

The first striking observation is the similarity of gene expression patterns among both variants of Signatures 3 and 13 and all other cloud signatures (2C and 8C). Since Signature 3 and 13 are considered to be associated with homologous recombination deficiency and APOBEC activity respectively, in what follows we refer to this group of signatures as HRD-APOBEC signature group. Note that Signature 2 is also known as an APOBEC related signature but the group includes only Signature 2C but not 2D. Below, we will discuss insights obtained for the age-related Signatures and the APOBEC signatures, and also provide independent supporting evidence from literature. Given expression correlation similarity within the members of the HRD-APOBEC group (all positively correlated with cell cycle, DNA repair, and immune response), we defer the analysis of this group to the next section where we look at this group through the lenses of mutated subnetworks.

#### The expression correlation analysis reveals important differences between the APOBEC signatures

Surprisingly, among 4 APOBEC related signatures (Signature 2C/D and 13C/D) Signature 2D has strikingly different correlation patterns compared to the remaining three APOBEC signatures. APOBEC activities are considered to be related to immune response. While the expression correlation patterns of all other APOBEC signatures are consistent with such understanding, Signature 2D exposure level has slightly negative correlation with immune response (2A, aC6). This is consistent with our previous observation that there is no positive correlation between Signature 2D and APOBEC expression [9].

In addition, Signature 2 exposure level is either not correlated (2D) or has a weak correlation (2C) with the cluster enriched with translesion synthesis (2, aC7 and mC4) whereas both Signature 13C and 13D show strong positive correlation. This last observation supports the previous claim that the difference between Signatures 2 and 13 is related to differences in the repair mechanism [34]. Specifically, it is suggested that mutations in Signatures 13 emerge when lesions created by APOBEC activity are repaired by DNA translesion polymerase, which inserts ‘C’ opposite to the damaged base while Signatures 2 occurs when the damaged base is simply paired with ‘A’.

#### Clock-like signatures 1D and 5D have different expression associations suggesting differences in their etiology

Although weaker than the correlation with the HRD-APOBEC Signature group, two clusters enriched in cell cycle function are positively correlated with Signature 1D (Fig. 2A, aC4 and aC5), which is consistent with the previous observation that Signature 1 is associated with aging [24], and thus postulated to be correlated with the number of cell divisions.

On the other hand, Signature 5D is negatively correlated with the expression of cell cycle genes although Signature 5 is also considered to be a clock-like signature. Instead, Signature 5D has a positive correlation with the cluster enriched in oxidative processes (Fig. 2A, aC1) and the cluster enriched in nucleotide excision repair (NER) pathway (Fig. 2B, mC1). The accumulation of oxidation base lesions is also assumed to be age-related [37], suggesting that Signature 5 might be related to oxidative damage. NER pathway is involved in neutralizing oxidative DNA damage [38] and Signature 5 has been also associated with smoking [39], which itself is associated with oxidative damage. Indeed Signature 5 was linked to the NER pathway in a recent study [40].

The positive correlation of Signature 1 with the expression of cell cycle genes and lack of such correlation for Signature 5 may explain the stronger association of Signature 5 with the age of patients than Signature 1 in breast cancer [9, 24] because cancer related cell division might obscure the association with a patient’s age.

### 3.2 Identifying mutated subnetworks associated with mutational signatures

The analysis of expression correlation clusters revealed different biological processes associated with some signatures but HR-APOBEC group have largely similar expression patterns. Complementary to the expression analysis, we next searched for possible associations with subnetworks of mutated genes. Some mutational signatures can arise due to endogenous causes; aberrations in genes responsible for different DNA repair mechanisms can lead to the malfunctioning of the corresponding repair process, leaving errors not repaired and in turn generating specific patterns of mutations. We applied NETPLHIX, a method to identify phenotype associated subnetworks, which can help to uncover a subnetwork of genes whose alterations are potentially causative of specific mutational signatures directly or indirectly. Note that not all mutational signatures have such association with mutated pathways. Mutational signatures arising from environmental exposure, age, or other external factors are not necessarily expected to have casual associations with mutated subnetworks.

Figure 3 shows all statistically significant subnetworks (phenotype permutation test; see Materials and Methods) identified by NETPHIX and their alteration profiles. See Materials and Methods (Section 5.2) for how the module for each signature was selected. The extended subnetworks obtained with less stringent cutoffs are shown in Figure S1.

As expected, no modules are found to be significantly associated with the age related signatures 1D and 5D. In addition, consistently with the previous studies that linked the genes underlying the HRD to Signature 3 in breast cancer [41], the subnetworks identified for Signature 3 C/D contain BRCA1 and BRCA2 genes, two important genes in HR-mediated double-strand break (DSB) repair. The agreement of the modules identified by NETPHIX with the current knowledge confirms its ability to correctly infer mutated subnetworks associations with signatures.

**Figure 3:**
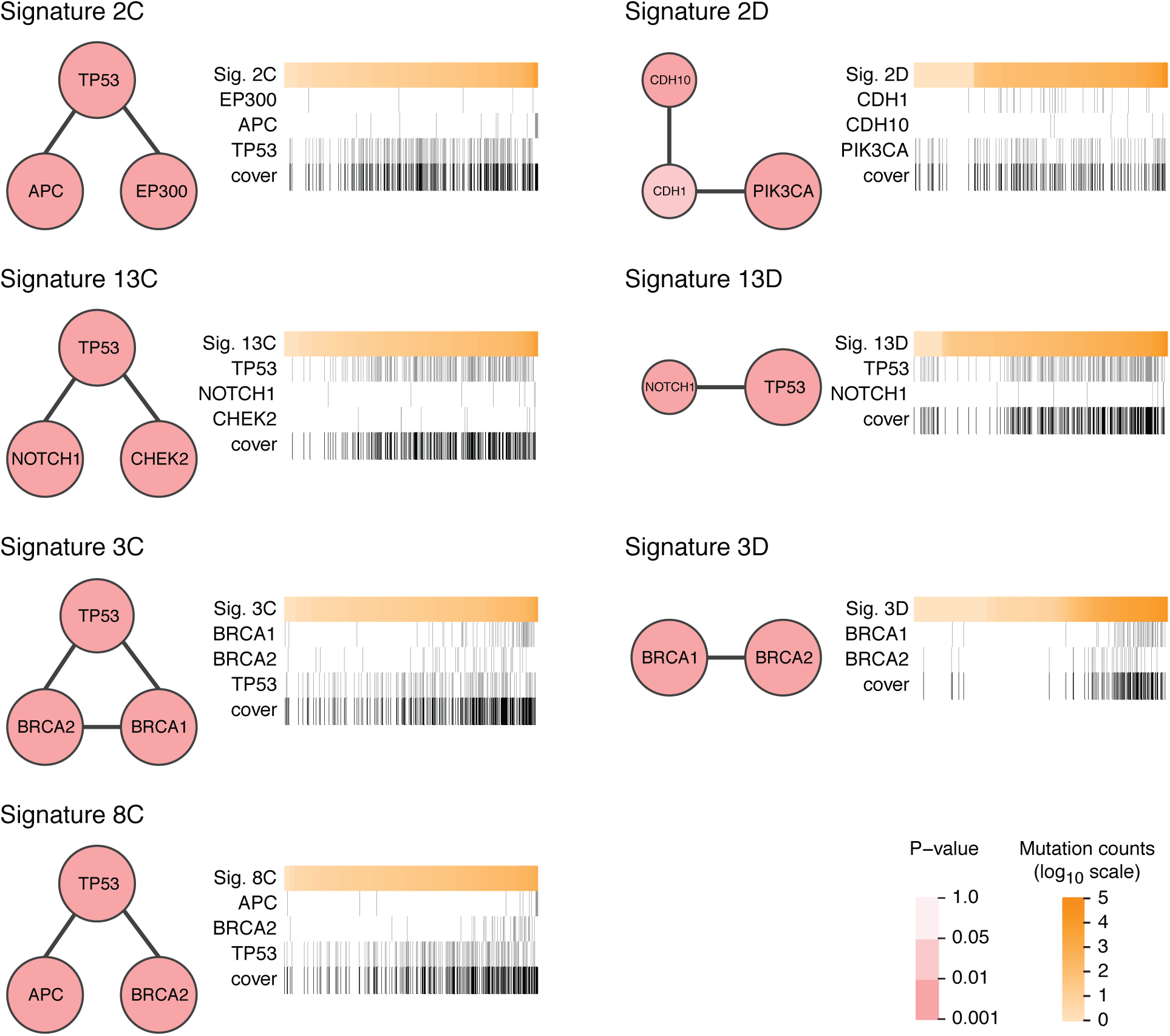
Subnetworks identified by NETPHIX for mutational Signatures 2C, 2D, 13C, 13D, 3C, 3D, and 8C. Panel for each signature consists of a network view of a module (left) and a heatmap showing an association of module gene alterations with signature strength across patients (right). The network node size indicates the gene robustness (regarding NETPHIX results for different random initialization runs of SIGMA) while the darkness of red color represents its individual association score (empirical p-value based on phenotype permutation test). Each heatmap shows the number of mutations attributed to a given signature for all patients (orange; top row; log10 scale) sorted from low to high (columns). For each gene in the module, gene alteration information observed in each patient are shown in gray, while patients not altered are in white. The last row shows the alteration profile of the entire subnetwork in black. Only subnetworks significant in phenotype associations are shown; results for signatures 1D and 5D were not significant.

Encouraged by the results, we examined the remaining subnetworks identified by NETPHIX. Among statistically significant modules, TP53 was included in all modules associated with cloud signatures. TP53 is known to play a crucial role in DNA damage responses, including DSB repair. We note that its dysfunction could contribute to increased mutation burden and in turn to the emergence of cloud mutations independently of mutagenic processes underlying individual signatures. However, whether or not TP53 mutations are causal or are a result of yet another mutagenic process cannot be concluded from this study. Complicating this picture, a recent study demonstrated that p53 controls expression of the DNA deaminase APOBEC3B suggesting a possible mechanism by which mutations in p53 can promote APOBEC expression [42] and thus APOBEC related mutations. Hence the reason for the strong association of TP53 with cloud mutational signatures require further investigation.

Compared to the modules obtained from expression analysis, the analysis with genetic alterations offers a better differentiation among the signatures in the HRD-APOBEC group. While most of the signatures in the group contain TP53, they include different genes in the modules. In the subnetworks associated with Signatures 13 C/D, TP53 is accompanied by NOTCH1; NOTCH pathway regulates many aspects of metazoan development including the control of proliferation and differentiation. CHEK2 is selected in addition to TP53 and NOTCH1 for Signature 13C. CHEK2 is a tumor suppressor regulating a cell cycle checkpoint and mutations in the gene confer an increased risk for breast cancer [43, 44]. CHEK2 plays multiple roles in DNA damage response [45], including DSB repair in the emergence of clustered APOBEC related mutations.

In the subnetwork associated with Signature 2C, TP53 is accompanied by APC (Adenomatous Polyposis Coli), which is a tumor-suppressor gene frequently mutated in Colorectal cancer (CRC) and involved in the Wnt signalling pathway. A recent study linked APC to several DNA repair mechanisms, including the base excision repair (BER) pathway [46], DSB repair [47] and genomic stability [48, 49].

Finally, the subnetwork for Signature 2D (dispersed, APOBEC related signature) consists of PIK3CA, CDH1 and CDH10 genes and is completely different from the subnetwork corresponding to the cloud variant of Signature 2 and other HR-APOBEC related signatures. Previous studies have found that some recurring mutations in PIK3CA are consistent with Signature 2 and may result from APOBEC activities [12, 50]. However, our analysis associated PIK3CA mutations with signature 2 even after removing point mutations attributed to Signature 2, suggesting a complex relation between Signature 2 and PIK3CA mutations.

In addition to PIK3CA the subnetwork associated with Signature 2 has two Cadherin genes: CDH1 and CDH10. Cadherins are important in maintenance of cell adhesion and polarity, and alterations of these functions can contribute to tumorigenesis. CDH1 germline mutations have been associated with hereditary lobular breast cancer [51], while a recent study linked mutations in CDH1 and PIK3CA mutations to the immune-related invasive lobular carcinoma of the breast [52]. In breast cancer, mutations in CDH1-PIK3CA module are mutually exclusive with mutations in TP53 and are strongly enriched in Luminal B subtype [53]. This, combined with the differences in expression correlations noted in the previous section, suggest that the etiology of Signature 2D is very different from the one of other APOBEC mutational signatures (Signature 2C and 13).

## 4 Conclusions

In this study, we used two different approaches to gain insights into the etiology of mutational processes in breast cancer. The first approach utilized gene expression data and identified gene modules whose expression correlates with mutation counts attributed to mutational signatures. The second approach focused on the identification of subnetworks of genes whose alterations are associated with each signature. The two approaches provided complementary insights.

The expression correlation based approach allowed us to uncover important differences between clock-like signatures 1 and 5. Clock-like signatures can occur from life long exposure to naturally occurring mutagenic processes, thus related to aging. The most prominent clock-like signatures are Signature 1 and 5. Signature 1, a relatively well characterized clock-like signature, is considered to be the result of an endogenous mutational process related to spontaneous deamination of 5-methylcytosine. Each cell division provides an opportunity for such mutations to occur. The etiology of Signature 5 is less clear. Our expression based analysis revealed that, differently from Signature 1, Signature 5 is not positively correlated with expression of cell cycle genes. Instead, we found an association of Signature 5 with oxidation process. This observation is consistent with several previous findings. Accumulation of oxidation base lesions is assumed to be related to aging [37] as well as smoking while the association of Signature 5 with smoking was observed from a previous study [39]. Another supporting evidence is provided by the association of Signature 5 with nucleotide excision repair (NER) pathway which was shown to be involved in neutralizing oxidative DNA damage [38].

While expression based analysis was very valuable for understanding the differences between Signatures 1 and 5, many signatures especially in the HRD-APOBEC signature group exhibit similar expression correlation patterns. The mutated pathway analysis better elucidated the differences among these signatures. In particular, Signatures 3D is associated with subnetwork consisting of BRCA 1/2 genes while the subnetwork associated with cloud of this signature (3C) contains TP53 additionally. The results of mutated subnetwork analysis reveals also the association of mutations in tumor suppressor APC for two different cloud signatures and NOTCH1 mutations for both variants of Signature 13.

In order to increase the probability that inferred mutated subnetworks are causal, we removed the mutations attributed to the signature of interest. This eliminates the possibility that the mutations resulted directly from the mutagenic process underlying the signature although it still does not guarantee causality. In particular, the consistent presence of TP53 in the subnetworks associated with cloud signatures makes it tempting to speculate that mutations in TP53 generally increase the mutation rates leading to an increase in cloud mutations. However, other indirect reasons for this association cannot be ruled out.

Our analysis also showed unique properties of Signature 2D relative to the remaining APOBEC signatures. This signature is the only signature associated with PIK3CA and not TP53. Previous studies have found that several recurring mutations in PIK3CA are consistent with Signature 2 [12, 50]. However, our analysis indicates that even after removing mutations attributed to Signature 2, the association between PIK3CA mutations and Signature 2D remains. Another known cancer gene present in this subnetwork is CHD1. CHD1 is linked to Hereditary Diffuse Gastric Cancer where it occur in about 40% of patients [54, 55] and to hereditary lobular breast cancer [56]. Invasive lobular carcinoma is characterized by a unique immune signature [57] which might provide additional insights to the etiology of Signature 2.

Taken together, this study demonstrates the utility of the two complementary approaches for studying mutational signatures in cancer and provided several new insights into the etiology of mutational signatures.

## 5. Materials and Methods

### 5.1 Data

We analyzed the gene expression data and the somatic mutations in the cohort of 560 breast cancer (BRCA) whole-genomes published by Nik-Zainal *et al.* [23]. We downloaded the normalized gene expression data for 266 BRCA patients from Table S7 in [23]. The mutation data were downloaded from ICGC data portal (https://dcc.icgc.org, release 22). The most likely assignments of mutational signatures to 3,479,652 individual point mutations were generated with SIGMA [9] using 12 COSMIC signatures identified as active in BRCA (Signatures 1, 2, 3, 5, 6, 8, 13, 17, 18, 20, 26, and 30; https://cancer.sanger.ac.uk/cosmic/signatures). To increase SIGMA’s robustness with respect to random initialization used in its learning process, we computed the majority assignments over 31 random initialization runs of SIGMA. In addition, we subdivided all mutations (and their associated signatures) into two groups – sequentially dependent close-by **C**loud (or clustered) mutations and independent **D**ispersed (or sky) mutations, as classified by SIGMA. For further analysis, we used only sufficiently abundant mutational signatures for cloud or dispersed mutations whose overall exposure levels are above 10% within each group. This created 10 different phenotype profiles for Signatures 1D, 2C/D, 3C/D, 5D, 8C/D, and 13C/D, where the numbering refers to the COSMIC signature index and C/D denotes signatures attributed to close-by cloud and dispersed mutations. For each patient, we computed signature profiles based on the patient mutation counts assigned to each specific signature, separating cloud and dispersed mutations. The mutational signature profiles were used as phenotype profiles in the expression correlation and mutated pathway analyses (Figure 1A).

For the gene level alteration information (the bottom matrix in Figure 1A), we utilized all somatic point mutations and small indels for the same 560 patients data. In processing the somatic mutation data, we defined a gene to be altered if it has at least one non-silent mutation in its genomic region. In addition to somatic mutations, DNA repair genes can undergo alternative mechanisms of inactivation including pathogenic germline variants and promoter hypermethylation. A recent paper highlighted the importance of these mechanisms in inactivating the homologous recombination pathway [2]. To account for these additional sources of inactivation, we also defined a gene to be altered in a patient if the gene is annotated as being biallelic inactivated in the patient as defined in Supplementary table 4a and 4b from [2]. The gene alteration information is used to find mutated subnetworks associated with each signature (Figure 1C). When computing association with a specific signature, we further refined the information to increase the likelihood that the association is causative. Specifically, the gene alteration information for the association analysis with a specific mutational signature were constructed after excluding the mutations attributed to the given mutational signature (we assumed that such mutations are the effects of the signature rather than the cause). The assignment of mutations to signatures was performed using SIGMA (see above). In addition, we removed all indels when we considered the associations with Signature 3 and 8 as these signatures are believed to be associated with HRD and high burden of indels.

### 5.2 Expression correlation analysis details

To identify expression based pathways that are associated with signatures, we used correlation analysis followed by clustering of correlation patterns. Specifically, we first computed Spearman correlation coefficient of the expression level and mutation count for each pair of genes and mutational signatures. We then selected the genes exhibiting significant correlation with at least one of 10 mutational signatures; the expression of a gene is considered significantly correlated with a signature if |*corr*| ≥0.3 and adjusted *pv ≤*0.005 (*corr* is Spearman correlation coefficient, BH corrected p-value). The procedure selected 3,763 genes. We then clustered the genes based on their correlation pattern using a consensus K-means algorithm; running K-means clustering 100 times with random start and varying *k* from 5 to 50 and subsequently running hierarchical clustering with consensus matrix from 100 runs of K-means. GO enrichment analysis was performed using hypergeometric test and significant terms were selected with nominal p-value *<* 0.05. The final 7 clusters and enrichment analysis results are summarized in Fig. 2A.

To take a closer look at DNA repair genes, we performed similar analysis with genes in GO DNA metabolic process. 184 genes are selected with the same significance cut-offs. The hierarchical clustering of the consensus clustering for 100 K-means (*k* = 2 to 20) generated 4 clusters shown in Fig. 2B. The enrichment analysis was performed using hypergeometric test with only the genes in GO DNA metabolic process as the background, and only for the GO terms with significant overlaps with GO DNA metabolic process (at least 2 genes in common and p-value of the intersection *<* 0.05).

### 5.3 Mutation analysis details

To find alteration based pathways for signatures, we adapted a recently developed method, NETPHIX which identifies mutated subnetworks associated with a continuous phenotype [36]. Given gene alteration information of cancer samples and continuous phenotype values for the same samples, NETPHIX aims to identify a connected subnetwork whose aggregated alterations are associated with the phenotype of interest (mutation counts for cancer signatures in this study). NETPHIX utilizes functional interaction information among genes and enforces the identified genes to be *connected* in the network while, at the same time, making sure that the aggregated alterations of these genes are significantly *associated* with the given phenotype. In addition, in its integer linear program formulation, NETPHIX recognizes that cancer driving mutations tend to be mutually exclusive [58, 59, 60, 53, 20, 61] and incorporates this property in its objective function [36].

For each mutational signature, we normalized mutation counts by taking log and subsequently computing z-scores. For functional interactions among genes, we used the data downloaded from STRING database version 10.0 (https://string-db.org), only including the edges with high confidence scores (≥900). Genes altered in less than 1% of patients were removed from further consideration. We ran NETPHIX for each mutational signature with density constraint of 0.5 and for a bounded size modules *m* from 1 to 7. The appropriate *m* was selected by examining the increase of the objective function values and the significance of the solution using permutation tests. Specifically, the best *m* was selected as maximal index for which the optimal objective function increased at least 5% with respect to previous index and the phenotype p-value did not increase, with this property holding for all smaller indices. The permutation test is computed by permuting the phenotype (the mutation counts for each signature in this case) and comparing the objective function value to the ones obtained with the permuted phenotypes. We define the identified module to be significant if the FDR adjusted *p*-value is less than 0.1 (see [36] for details).

## Acknowledgements

This research was supported in part by the Intramural Research Programs of the National Library of Medicine at National Institutes of Health, USA. RS was supported by Len Blavatnik and the Blavatnik Family foundation. FV was supported, in part, by the University of Padova grants “SID2017” and “STARS: Algorithms for Inferential Data Mining”.

## Supplementary Materials

**Figure S1:**
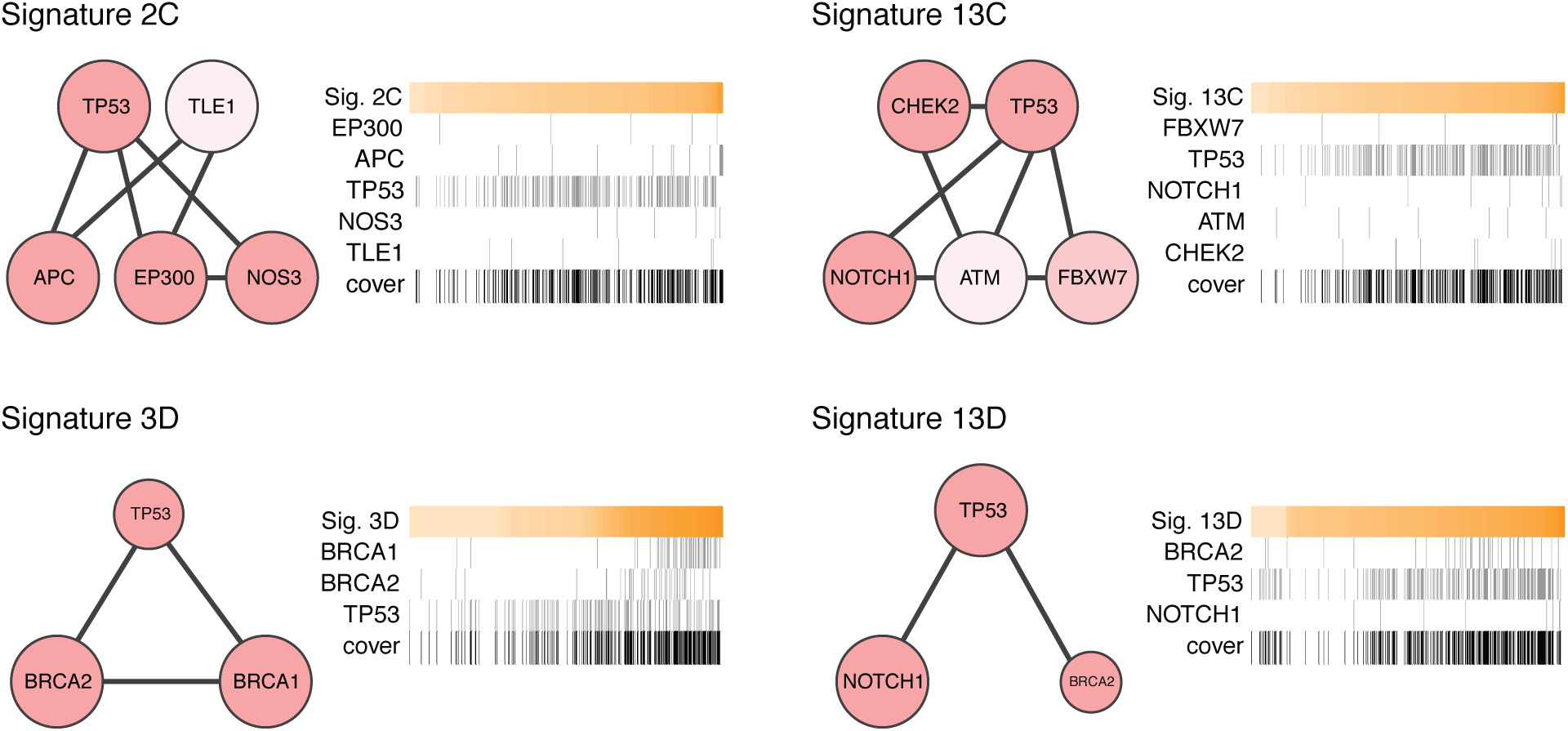
Subnetworks identified by NETPHIX for mutational Signatures 2C, 13C, 13D, and 3D using less stringent cut-off (refers to Figure 3). The best *m* (the module size) using less stringent cut-offs was selected as maximal index for which the optimal objective function increased at least 1% with respect to previous index and the phenotype p-value did not increase. Panel for each signature consists of a network view of a module (left) and a heatmap showing the association of module gene alterations with signature strength across patients (right). The network node size indicates the gene robustness (regarding NETPHIX results for different random initialization runs of SIGMA) while the darkness of red color represents its individual association score (*p*-value). Each heatmap shows the number of mutations attributed to a given signature for all samples (orange; top row; log10 scale) sorted from low to high (columns). For each gene in the module, gene mutations observed in each sample caused by other signatures are shown in gray, while samples not altered are in white. The last row shows the mutation profile of the entire subnetwork in black. Only subnetworks that changed with respect to the normal cut-offs (see Figure 3 and Materials and Methods) are shown. Results for Signatures 2D, 3C and 8C did not change with respect to the normal cut-offs and results for Signatures 1D and 5D stayed insignificant (the FDR adjusted *p*-value above than 0.1).

**Table S1:**
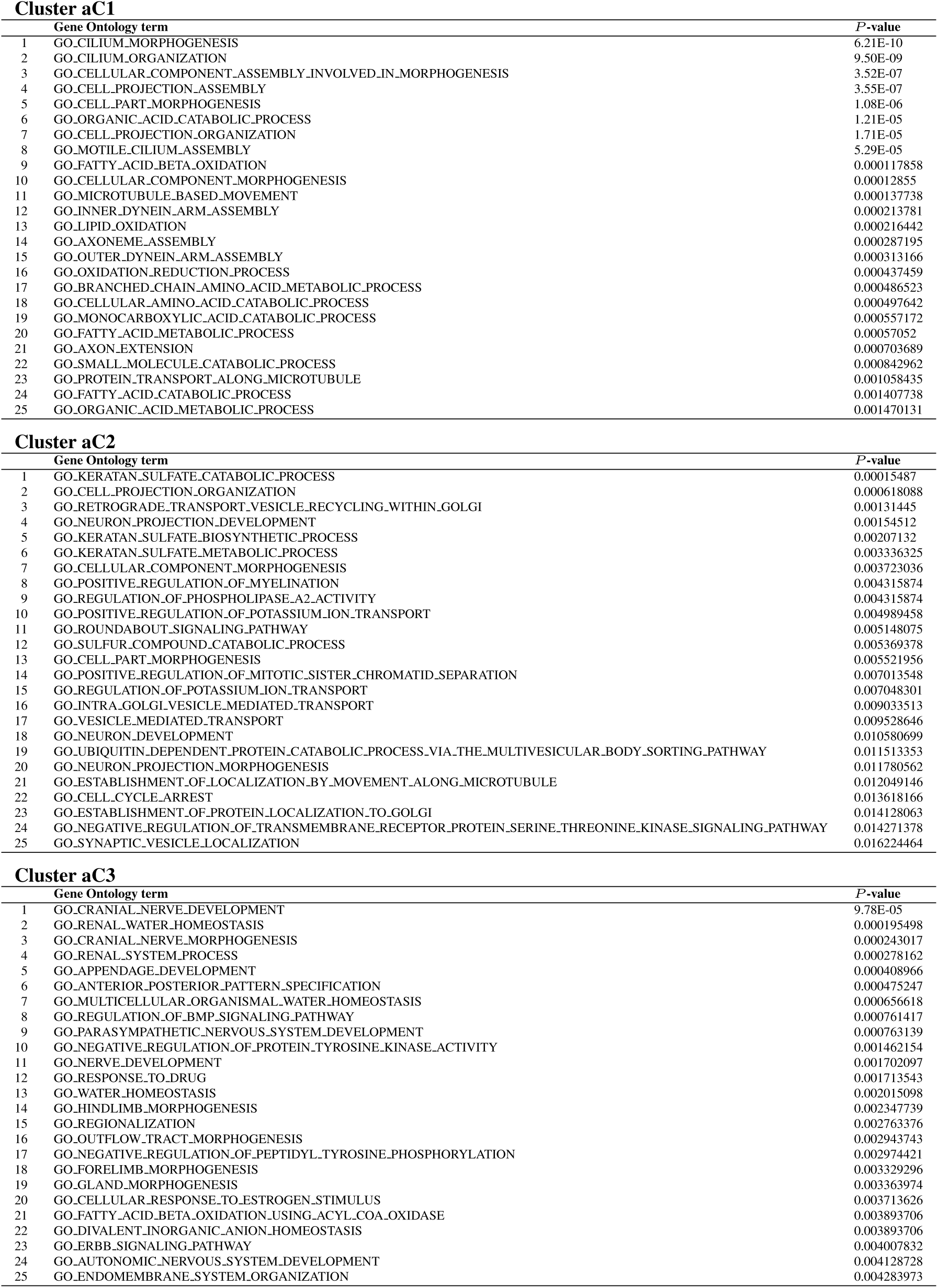
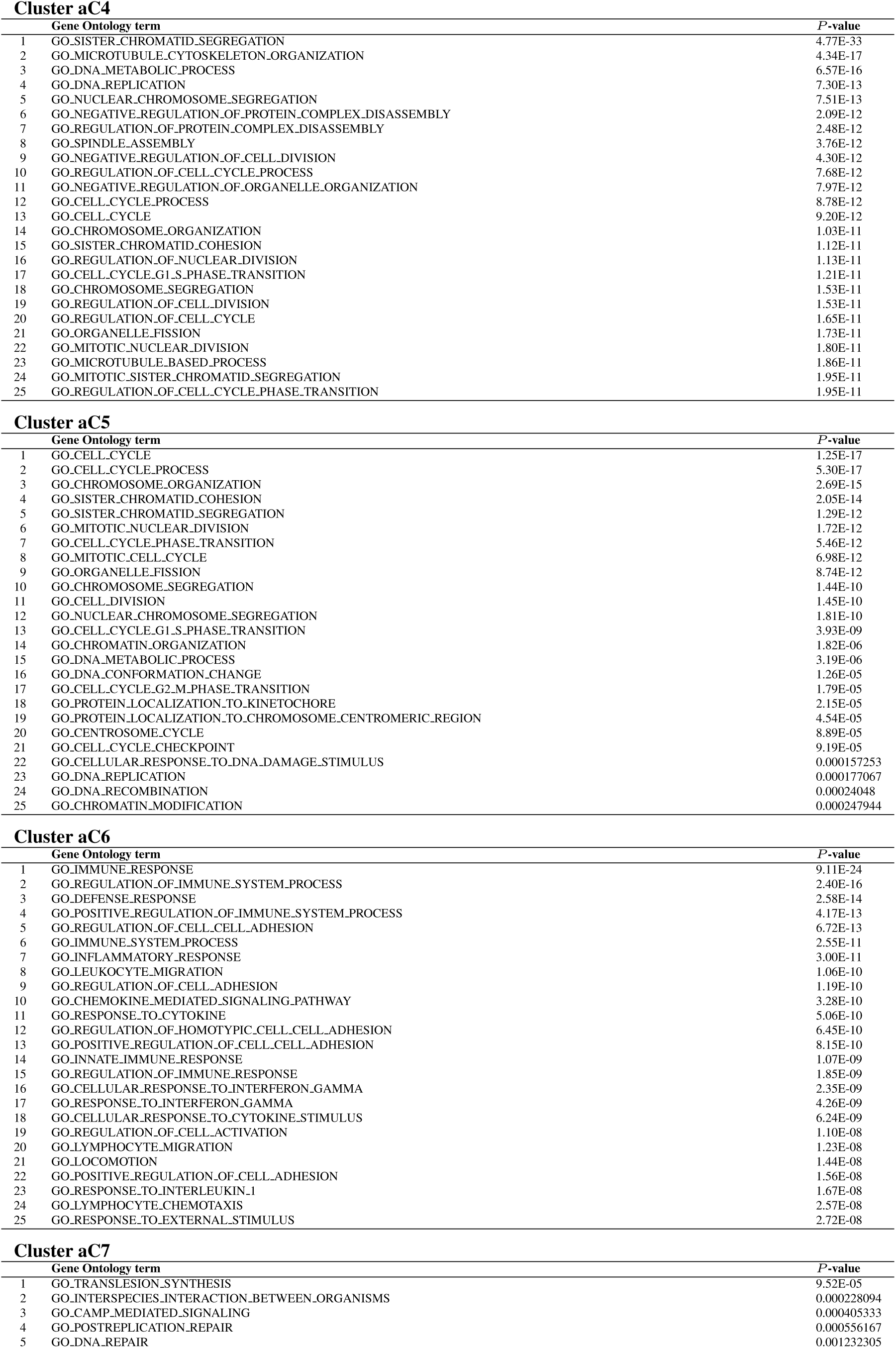
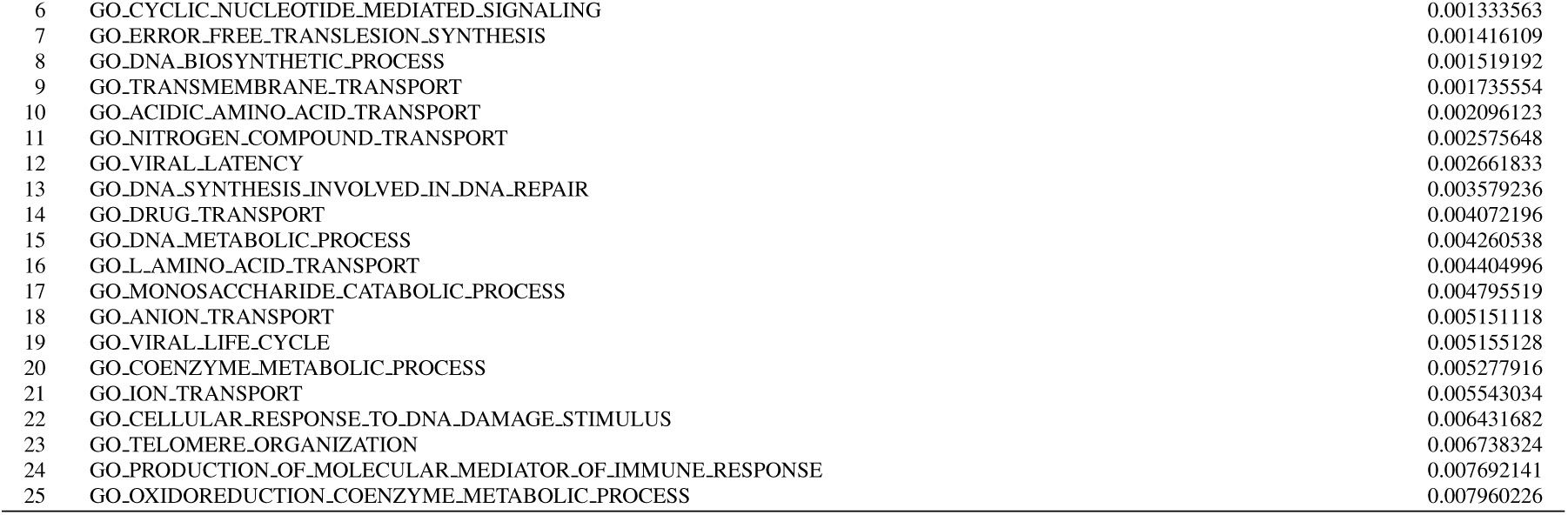
Gene Ontology enrichment analysis (refers to Figure 2A). Gene Ontology terms enriched in 7 gene expression correlation modules for all genes significantly correlated with at least one signature (refers to Figure 2A). Top 25 GO terms with *p*-value above 0.05 are shown for each cluster/module.

**Table S2:** Gene Ontology enrichment analysis (refers to Figure 2B). GO terms enriched in 4 gene expression correlation modules for DNA metabolic process genes significantly correlated with at least one signature. Top 25 GO terms with *p*-value above 0.05 are shown for each cluster/module.

| Cluster mC1 |  |  |
| --- | --- | --- |
|  | Gene Ontology term | P-value |
| 1 | GO_NUCLEOTIDE_EXCISION_REPAIR | 0.008852967 |
| 2 | GO_CELLULAR_RESPONSE_TO_LIGHT_STIMULUS | 0.031320019 |
| 3 | GO_NUCLEOTIDE_EXCISION_REPAIR_PREINCISION_COMPLEX_ASSEMBLY | 0.039676567 |
| 4 | GO_POSITIVE_REGULATION_OF_PROTEIN_MODIFICATION_PROCESS | 0.039809348 |
| 5 | GO_MISMATCH_REPAIR | 0.04920754 |
| Cluster mC2 |  |  |
|  | Gene Ontology term | P-value |
| 1 | GO_CELL_CYCLE_PROCESS | 9.42E-08 |
| 2 | GO_CELL_CYCLE | 1.98E-07 |
| 3 | GO_MITOTIC_CELL_CYCLE | 4.49E-07 |
| 4 | GO_CELL_CYCLE_PHASE_TRANSITION | 3.65E-06 |
| 5 | GO_CELL_CYCLE_G1_S_PHASE_TRANSITION | 4.86E-06 |
| 6 | GO_DNA_REPLICATION_INITIATION | 1.38E-05 |
| 7 | GO_REGULATION_OF_DOUBLE_STRAND_BREAK_REPAIR_VIA_HOMOLOGOUS_RECOMBINATION | 0.000477316 |
| 8 | GO_DNA_DEPENDENT_DNA_REPLICATION | 0.000683233 |
| 9 | GO_DNA_REPLICATION | 0.000871556 |
| 10 | GO_RESPONSE_TO_VITAMIN | 0.001009416 |
| 11 | GO_REGULATION_OF_DOUBLE_STRAND_BREAK_REPAIR | 0.00318849 |
| 12 | GO_DNA_GEOMETRIC_CHANGE | 0.004523221 |
| 13 | GO_REGULATION_OF_TRANSCRIPTION_INVOLVED_IN_G1_S_TRANSITION_OF_MITOTIC_CELL_CYCLE | 0.004820393 |
| 14 | GO_MITOTIC_G2_M_TRANSITION_CHECKPOINT | 0.004820393 |
| 15 | GO_NEGATIVE_REGULATION_OF_CELL_CYCLE_G2_M_PHASE_TRANSITION | 0.00646435 |
| 16 | GO_RESPONSE_TO_ORGANIC_CYCLIC_COMPOUND | 0.006515012 |
| 17 | GO_REGULATION_OF_CELL_CYCLE_G2_M_PHASE_TRANSITION | 0.006574288 |
| 18 | GO_NEGATIVE_REGULATION_OF_CELL_CYCLE_PROCESS | 0.009287607 |
| 19 | GO_REGULATION_OF_MITOTIC_CELL_CYCLE | 0.009801488 |
| 20 | GO_NEGATIVE_REGULATION_OF_MITOTIC_CELL_CYCLE | 0.010394224 |
| 21 | GO_REGULATION_OF_DNA_RECOMBINATION | 0.010620267 |
| 22 | GO_REGULATION_OF_CELL_CYCLE_PHASE_TRANSITION | 0.010990845 |
| 23 | GO_REGULATION_OF_CELL_CYCLE_PROCESS | 0.012683361 |
| 24 | GO_STRAND_DISPLACEMENT | 0.013779842 |
| 25 | GO_RESPONSE_TO_ALCOHOL | 0.014076202 |
| Cluster mC3 |  |  |
|  | Gene Ontology term | P-value |
| 1 | GO_PROTEIN_MONOUBIQUITINATION | 0.000207747 |
| 2 | GO_CELL_CYCLE | 0.000244027 |
| 3 | GO_MITOTIC_CELL_CYCLE | 0.000565274 |
| 4 | GO_CELL_DIVISION | 0.002250474 |
| 5 | GO_REGULATION_OF_CIRCADIAN_RHYTHM | 0.003484062 |
| 6 | GO_CELL_CYCLE_CHECKPOINT | 0.004693563 |
| 7 | GO_MITOTIC_NUCLEAR_DIVISION | 0.005030615 |
| 8 | GO_RESPONSE_TO_DRUG | 0.005142209 |
| 9 | GO_CELL_CYCLE_PHASE_TRANSITION | 0.008091532 |
| 10 | GO_CELL_CYCLE_PROCESS | 0.008940729 |
| 11 | GO_METHYLATION_DEPENDENT_CHROMATIN_SILENCING | 0.00952548 |
| 12 | GO_DNA_RECOMBINATION | 0.009605795 |
| 13 | GO_REPRODUCTION | 0.009964206 |
| 14 | GO_ORGANELLE_FISSION | 0.015820914 |
| 15 | GO_MICROTUBULE_ORGANIZING_CENTER_ORGANIZATION | 0.016457806 |
| 16 | GO_CELL_CYCLE_G1_S_PHASE_TRANSITION | 0.016483658 |
| 17 | GO_MITOTIC_CELL_CYCLE_CHECKPOINT | 0.018392498 |
| 18 | GO_REGULATION_OF_DNA_DEPENDENT_DNA_REPLICATION | 0.018861964 |
| 19 | GO_CELL_DEATH | 0.024499591 |
| 20 | GO_RHYTHMIC_PROCESS | 0.025059498 |
| 21 | GO_NEGATIVE_REGULATION_OF_CELL_DEATH | 0.025275716 |
| 22 | GO_MITOTIC_RECOMBINATION | 0.026039036 |
| 23 | GO_G1_DNA_DAMAGE_CHECKPOINT | 0.028636609 |
| 24 | GO_PROTEIN_UBIQUITINATION_INVOLVED_IN_UBIQUITIN_DEPENDENT_PROTEIN_CATABOLIC_PROCESS | 0.028636609 |
| 25 | GO_SISTER_CHROMATID_SEGREGATION | 0.028636609 |

Cluster mC4
|  | Gene Ontology term | P-value |
| --- | --- | --- |
| 1 | GO_REGULATION_OF_SYMBIOSIS_ENCOMPASSING_MUTUALISM_THROUGH_PARASITISM | 0.000293075 |
| 2 | GO_REGULATION_OF_VIRAL_GENOME_REPLICATION | 0.000823086 |
| 3 | GO_NEGATIVE_REGULATION_OF_VIRAL_GENOME_REPLICATION | 0.000841816 |
| 4 | GO_REGULATION_OF_MULTI_ORGANISM_PROCESS | 0.001610789 |
| 5 | GO_NEGATIVE_REGULATION_OF_VIRAL_PROCESS | 0.004263298 |
| 6 | GO_NEGATIVE_REGULATION_OF_MULTI_ORGANISM_PROCESS | 0.005827031 |
| 7 | GO_POSITIVE_REGULATION_OF_VIRAL_PROCESS | 0.006665488 |
| 8 | GO_NEGATIVE_REGULATION_OF_GENE_SILENCING | 0.007951779 |
| 9 | GO_POSITIVE_REGULATION_OF_MULTI_ORGANISM_PROCESS | 0.008344127 |
| 10 | GO_IMMUNE_EFFECTOR_PROCESS | 0.01134109 |
| 11 | GO_ERROR_FREE_TRANSLESION_SYNTHESIS | 0.012875934 |
| 12 | GO_VIRAL_LATENCY | 0.013134845 |
| 13 | GO_REGULATION_OF_CELLULAR_SENESCENCE | 0.013134845 |
| 14 | GO_ANATOMICAL_STRUCTURE_HOMEOSTASIS | 0.014981217 |
| 15 | GO_DNA_DEPENDENT_DNA_REPLICATION | 0.016367536 |
| 16 | GO_RETROGRADE_PROTEIN_TRANSPORT_ER_TO_CYTOSOL | 0.01756138 |
| 17 | GO_TELOMERE_ORGANIZATION | 0.019277733 |
| 18 | GO_REGULATION_OF_GENE_SILENCING | 0.019478047 |
| 19 | GO_PYRIMIDINE_RIBONUCLEOSIDE_CATABOLIC_PROCESS | 0.019840394 |
| 20 | GO_SINGLE_ORGANISM_BIOSYNTHETIC_PROCESS | 0.024977871 |
| 21 | GO_BLASTOCYST_DEVELOPMENT | 0.025010683 |
| 22 | GO_DNA_STRAND_ELONGATION | 0.025237984 |
| 23 | GO_PYRIMIDINE_RIBONUCLEOSIDE_METABOLIC_PROCESS | 0.028101631 |
| 24 | GO_RIBONUCLEOSIDE_CATABOLIC_PROCESS | 0.028101631 |
| 25 | GO_REGULATION_OF_CELL_AGING | 0.028101631 |

